# Tracking transitional probabilities and segmenting auditory sequences are dissociable processes in adults and neonates

**DOI:** 10.1101/2021.09.02.458702

**Authors:** Lucas Benjamin, Ana Fló, Marie Palu, Shruit Naik, Lucia Melloni, Ghislaine Dehane-Lambertz

**Affiliations:** Cognitive Neuroimaging Unit, CNRS ERL 9003, INSERM U992, CEA, Université Paris-Saclay, NeuroSpin center, 91191 Gif/Yvette, France; Department of Neuroscience, Max Planck Institute for Empirical Aesthetics, 60322, Frankfurt a.M, Germany; Department of Neurology, NYU Grossman School of Medicine, 10016, New York, US

## Abstract

Since speech is a continuous stream with no systematic boundaries between words, how do pre-verbal infants manage to discover words? A proposed solution is that they might use the transitional probability between adjacent syllables, which drops at word boundaries. Here, we tested the limits of this mechanism by increasing the size of the word-unit to 4 syllables, and its automaticity by testing asleep neonates. Using markers of statistical learning in neonates’ EEG, compared to adult’ behavioral performances in the same task, we confirmed that statistical learning is automatic enough to be efficient even in sleeping neonates. But we also revealed that: 1) Successfully tracking transition probabilities in a sequence is not sufficient to segment it 2) Prosodic cues, as subtle as subliminal pauses, enable to recover segmenting capacities 3) Adults’ and neonates’ capacities are remarkably similar despite the difference of maturation and expertise. Finally, we observed that learning increased the similarity of neural responses across infants, providing a new neural marker to monitor learning. Thus, from birth, infants are equipped with adult-like tools, allowing to extract small coherent word-like units within auditory streams, based on the combination of statistical analyses and prosodic cues.

## Introduction

One of the main challenge encountered by infants to learn their native language and construct their lexicon is that words are rarely produced in isolation. Instead, words are embeded into multi word sentences with no silence or clear acoustic boundaries between them. Although word endings is sometimes signaled by subtle acoustical markers such as the lengthening of the last syllable, pitch change, slowing-down of the syllabic rate and less coarticulation between syllables, adults have usually difficulties to correctly segment sentences in words in an unknown natural language (***Wakefield et al., 1974***). However, when the experimental task is simplified and an artificial stream of concatenated words is created, they are able to use acoustical cues to discover the possible words as shown by their above chance accuracy in forced-choice tasks (***Bagou and Frauenfelder, 2018***). Similarly, neonates have been shown to detect these subtle variations in a binary situation in which they have to discriminate pseudo-words constituted of syllables either coming from inside a word (e.g. mati - */mati/* from mathematicien)or from two successive words (e.g. maty - */mati/* from panorama typique) (***Christophe et al., 1994***). However, the relative weights of these markers vary across languages (***Ordin et al., 2017***) and also within a language (i.e. they depend on the position of the word in the sentence and interact with other prosodic features such as lexical stress). Therefore, the robustness of these word-boundary cues is commonly estimated as insufficient for infants to segment natural speech in successive word units.

A second mechanism, based on the analysis of the transitions between syllables, has thus been proposed. Within a word, syllables have a fixed order, whereas any syllable can follow the last syllable of a word. Thus, a word boundary corresponds to a drop in the transition probabilities (TP) between consecutive syllables. To prove that the concept could apply for word learning in infancy, ***Saffran et al.*** (***1996a***) used a mini language of 4 tri-syllabic words and tested 8-month-old infants who listened for 3 mins to a continuous stream in which these words were concatenated with a flat intonation. The authors reported that infants were subsequently able to distinguish two different lists of isolated tri-syllables pseudo-words: one corresponding to the words (i.e. ABC: TP were equal to 1 between each syllable) and the other to PartWords formed by the two last syllables of a word and the first syllable of the next word for example (i.e. BCA’: TP were equal to 1 and 0.33). This result has been replicated multiple times (***Black and Bergmann, 2017***) and extended to non-linguistic stimuli (***Saffran et al., 1999***; ***Schön et al., 2008***) and to the visual domain (***Fiser and Aslin, 2002***). Sensitivity to statistics in sequences is also observed in animals (***Hauser et al., 2001***; ***Toro and Trobalón, 2005***; ***James et al., 2020***) indicating that the capacity of extracting transitional probabilities between subsequent elements is a robust general mechanism. It has been reported in asleep neonates (***Teinonen et al., 2009***; ***Fló et al., 2019***, ***2021a***); and also, to some extent, in inattentive adults (***Batterink and Choi, 2021***; ***Benjamin et al., 2021***; ***Toro et al., 2005***). Yet the limits of this learning mechanism are still to be determined such as the influence of development and expertise on the performances.

One of the limitations of statistical learning, already reported in the literature, is its interaction with alternative segmentation cues (***Black and Bergmann, 2017***) and especially its embedding in prosodic units. A word cannot straddle a prosodic boundary. Therefore, even if two syllables are always presented in succession, they are not attributed to the same word if a prosodic boundary separates them. This property is observed in adults (***Shukla et al., 2007***) and in 5 to 8-month-old babies (***Johnson and Tyler, 2010***; ***Shukla et al., 2011***). This result should not be surprising given the importance of prosody to structure the speech signal. A hierarchy of prosodic units (***Nespor and Vogel, 2006***) roughly parallel to the syntactic tree is used to improve speech comprehension in adults and to favor language acquisition in infants. For example, even at two-months of age, infants memorize better the phonetic content of a sentence with a well-formed prosodic contour relative to a word-list (***Mandel et al., 1994***). This advantage can be explained because statistical computations are limited to a few elements within the prosodic unit, relieving memory. Prosodic units also provide perceptual anchors, that help infants to note the reproducible location of some words at their edges, such as articles or proper name. Finally, the higher frequency of function words relative to content words has also been proposed as anchors favoring word discovery (***Hochmann et al., 2010***). To succeed in the complex task of constructing a lexicon from natural speech, infants have a toolbox of procedures at their disposal, whose relative contributions are important to understand but are currently underspecified.

We investigated here another putative limitation of statistical learning, which is the size of the words that can be learned. In fact, most, if not all, studies in infants have used tri-syllabic words. Is it due to particular experimental choices or is it because there is a hard limit to statistical computation especially in immature infants ? If it is the case, can subtle prosodic cues rescue word learning by segmenting the stream, allowing memory processes to deploy (***Fló et al., 2021a***)? To investigate these questions, we created a first artificial stream consisting of four quadri-syllabic words, pseudo-randomly concatenated without any prosodic cue, and a second one strictly identical to the first one but with a 25ms pause between each words, every four syllables. In previous artificial language studies using this short pause, adults reported not to notice it and were at chance when they had to choose which of the two streams had pauses (***Peña et al., 2002***). Nevertheless, pauses significantly improved their performances (***Peña et al., 2002***; ***Buiatti et al., 2009***). This pause was probably perceived as a vowel lengthening, which is a universal ending cue, for words but also for musical segments (***Tyler and Cutler, 2009***). In adults, final syllable lengthening improved tri-syllabic word segmentation (***Saffran et al., 1996b***). The authors proposed a putative order in the use of these cues, i.e. infants first rely on transitional probabilities, then notice that syllable lengthening coincides with a word ending to finally learn this new cue. Yet, this hypothesis remains untested, because the relative contribution of transitional probabilities and of this subtle prosodic cue was not assessed in this study.

We used high-density EEG (128 channels) to evaluate segmentation processes in neonates. EEG allows not only to observe different responses to test-words after learning, but also to track the learning trajectory while neonates are listening to the artificial stream. As the syllables have exactly the same length, their perception creates a regular evoked response, which is observed as a power increase at the frequency of the syllable presentation. If the syllables are perceived grouped in a quadri-syllabic word, the power should increase at ¼ of the syllable frequency (1/3 if tri-syllabic words are used). Such a power increase at the word frequency has indeed been reported for tri-syllabic words in adults (***Buiatti et al., 2009***; ***Batterink and Choi, 2021***; ***Benjamin et al., 2021***) and in 8-month-old infants (***Kabdebon et al., 2015***). We also presented to the neonates streams with pseudo-randomly concatenated syllables, with and without a pause every four syllables, to control whether the pause by itselfwas sufficientto induce a 4-syllable-rythm. In adults, inserting such a pause in a random stream did not produce any increase of power at the pause frequency (***Buiatti et al., 2009***). Therefore, in case of successful segmentation, we expected a significant power increase at the word frequency in the structured stream relative to the random stream. No changes, or perhaps a decrease, was expected at the syllabic rate, in line with previous reports in adults in which, perceiving the word induced a decrease of the entrainment at the syllabic rate (***Batterink and Choi, 2021***; ***Benjamin et al., 2021***).

After the learning phase, three types of test-words were presented in isolation: Words, PartWords and ShuffleWords (fig1). Successful word segmentation is commonly revealed by a significant difference between the measured response to Words and to PartWords. In Words, all transitional probabilities between syllables equal 1, while in PartWords (straddling two words), there is a drop in transitional probabilities that indicates an ill-formed word. In ShuffleWords, the two middle syllables of a Word were inverted, violating local position. Thus, while all the transitional probabilities were zero, all syllables were always heard in close proximity during the learning stream. This temporal proximity might induce memory errors and a false recognition of ShuffleWords as possible words. Indeed in longer words of six-syllables, neonates are not able to detect a shuffle of the middle syllables whereas they detect a shuffle of the edges syllables (***Ferry et al., 2016***).

**Figure 1.**
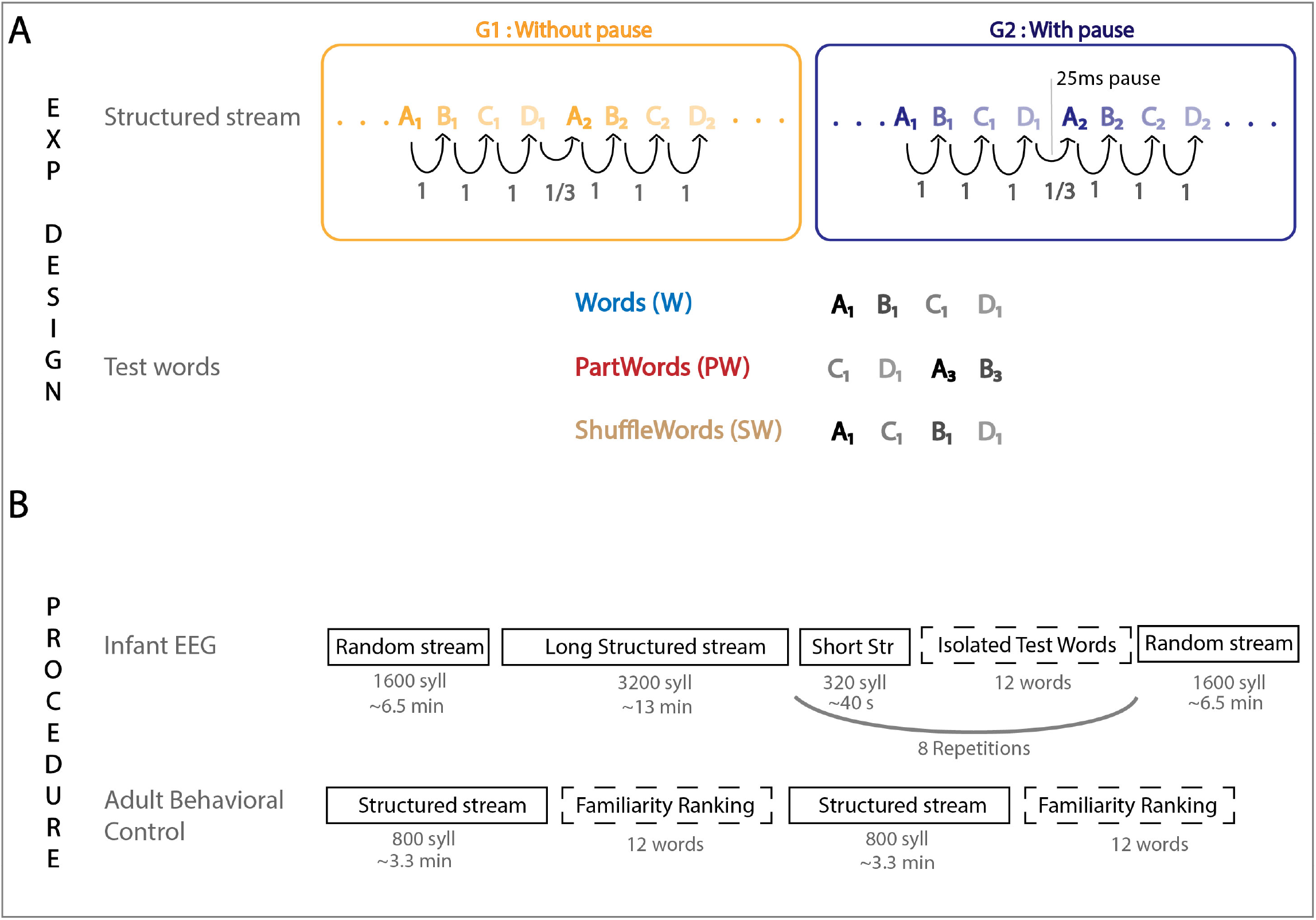
A: Experimental Design: Participants were first exposed to a structured stream comprising four quadri-syllabic words (called ABCD) then presented with 3 types of isolated test words. B: Experimental procedure: Neonates were tested asleep using high-density EEG (128 channels) while they were listening to random stream, structured stream, and isolated words. Adults were tested on a web platform. After familiarisation with the structured stream ( 3.3 mins), adults were asked to rank the familiarity on a scale (1 to 6). To avoid a bias to quadri-syllabic words, they were also presented with three other types of bi-syllabic test words (see methods for more details). Neonates were passively exposed to a structured stream (13 mins) followed by isolated test words.

Thus, our experimental design provided us with several markers of transitional probability computation and word segmentation that might be differently associated, opening the possibility to disentangle several steps or hypotheses of this classical learning task.(*H1: TP computation*) If infants computed TP and memorized the TP matrix, they should reject Words from ShuffleWords (1+1+1 vs 0+0+0) and eventually Words from PartWords (1+1+1 vs 1+0.33+1). (*H2: segmentation*) Stream segmentation should create an increase of neural entrainment at the word frequency. (*H3: complete memorization of the word*) should create a difference between words on one side, PartWords and ShuffleWords on the other side. (*H4: memory errors*) If Words are segmented and swap errors occur, ShuffleWords should not differ from Words due to the temporal proximity of the syllables belonging to the same Word. (*H5: first syllable memorization only*). This hypothesis could explain why words are preferred over PartWords in many statistical learning studies. As the typical trisylabic paradigm compare Words (ABC) to PartWords (BCA’), the difference observed could result from the encoding of the first syllable only (A vs B). In a recent study with tri-syllabic words, we indeed observed an ERP difference between words and PartWords for the first syllable, whereas no difference was recorded when the last syllable was incorrect (***Fló et al., 2021a***).

Finally, the comparison between the two groups of neonates, one listening to the stream without pauses and the other to the stream with pauses, should clarify the relative contribution of prosodic cues and transitional probabilities to word segmentation. Thereby advancing our understanding whether pauses rescue segmentation favoring the computation of transitional probabilities on smaller segments, or whether they are rather two independent mechanisms.

Neonates are two-decades far from a mature state in terms of linguistic abilities but also in terms of memory capacities. To our knowledge, no adult equivalent of the paradigm proposed here is available. Thus, we collected adult behavioral data as a mature model of the mechanisms we explored in neonates. We adapted the paradigm to make it shorter and to collect behavioral responses on a web-based procedure (see fig 1). Their results were surprisingly similar to those in infants, revealing common basic processes with similar limitations.

## Results

### Adults

Two groups of adults were tested online on a web platform. We analyzed the responses by items in a linear mixed-effects model in each group with FDR correction. For the stream without subliminal pause at the end of the word, Words and PartWords were similarly rated (p=0.26) and estimated more familiar than the ShuffleWords (W vs SW p<0.001, PW vs SW p<0.01). When subliminal pauses were added at the end of the words in the stream, all types of words were ranked differently (all p<0.001)with the following order: Words were judged more familiar than PartWords itself more familiar than ShuffleWords (Fig 1 A&B). To better visualize the difference in segmentation performances between the two groups, we calculated the difference in mean familiarity ranking given by each participant to Words and PartWords, and performed an unpaired unidirectional, t-test t(41)= 2.3, p=0.0128. The segmentation effect, seen as a positive value on the fig 2C, was larger when subliminal pauses were present (Fig 2 C).

**Figure 2.**
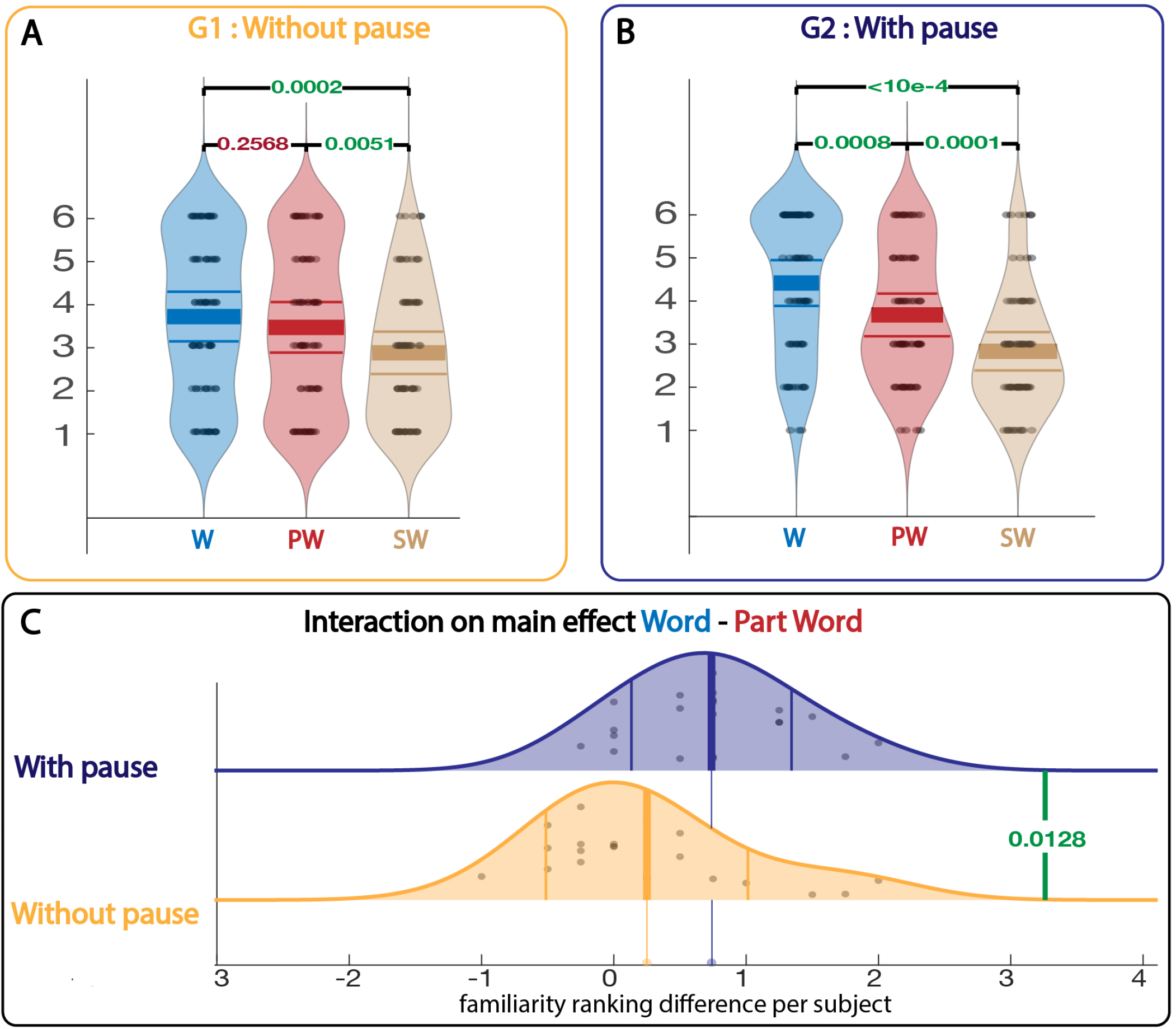
A,B: Results ofthe familiarity ranking tests by adults subjects. Here are represented the mean familiarity rank for each item. C: Interaction at the subject level between both groups on the main effect of segmenting (Word - PartWord). Dots represent familiarity ranking difference between Words and PartWords for each subject. p-value is estimated using one-way unpaired t-test.

Thus adults were able to distinguish Words from PartWords, indicating that they had correctly segmented the stream only if helped by a subliminal acoustic cue. Yet even when there was no pause, they rejected ShuffleWords because of null transitional probabilities.

### Infant EEG Experiment

EEG was recorded in two groups of healthy full-term neonates while they were listening to streams without pauses for the first group and with 25 ms pauses every four syllables for the second group. For each group, neonates were exposed first to 6.7 minutes (6.8 minutes for the group with pauses) of a stream in which syllables were randomly concatenated with a flat TP of 0.33, followed by 13.3 (resp. 13.7) minutes of the word-structured stream. After the exposure learning phase, a test phase followed in which they were exposed to isolated quadri-syllables sequences (Words, PartWords and ShuffleWords). To avoid interference with learning in the testing phase and to reinforce learning the structured materials, 40s-short segments of the structured stream were presented every 12 words during this phase. Finally, another 6.7 (resp 6.8) minutes of the random stream was presented to avoid a time confound in the comparison between random and structured streams.

#### Neural entrainment

As described in other studies on neural entrainment (***Kabdebon et al., 2015***; ***Buiatti et al., 2009***; ***Fló et al., 2021a***), there was a significant increase in power and Phase Locking Value (PLV) at the frequency of the syllables presentation compared to neighboring frequencies in both groups (with and without pause) and stream types (random and structured) (all ps<0.05 FDR corrected). No interaction was observed between groups and streams indicating a similar signal-to-noise ratio and comparable experimental conditions in the two groups (all ps>0.05 FDR corrected).

Stream segmentation should be revealed by a significant increase of power and/or phase locking value at the frequency of the words, i.e. ¼ of the syllabic frequency, relative to the random stream. In the first group (without pause), we failed to find this result, contrary to the second group (with pause), in whom a significant increase of both power and PLV was observed in several electrodes. Finally, the interaction between groups and streams was significant for both power and PLV on several electrodes (see fig3 all ps<0.05).

**Figure 3.**
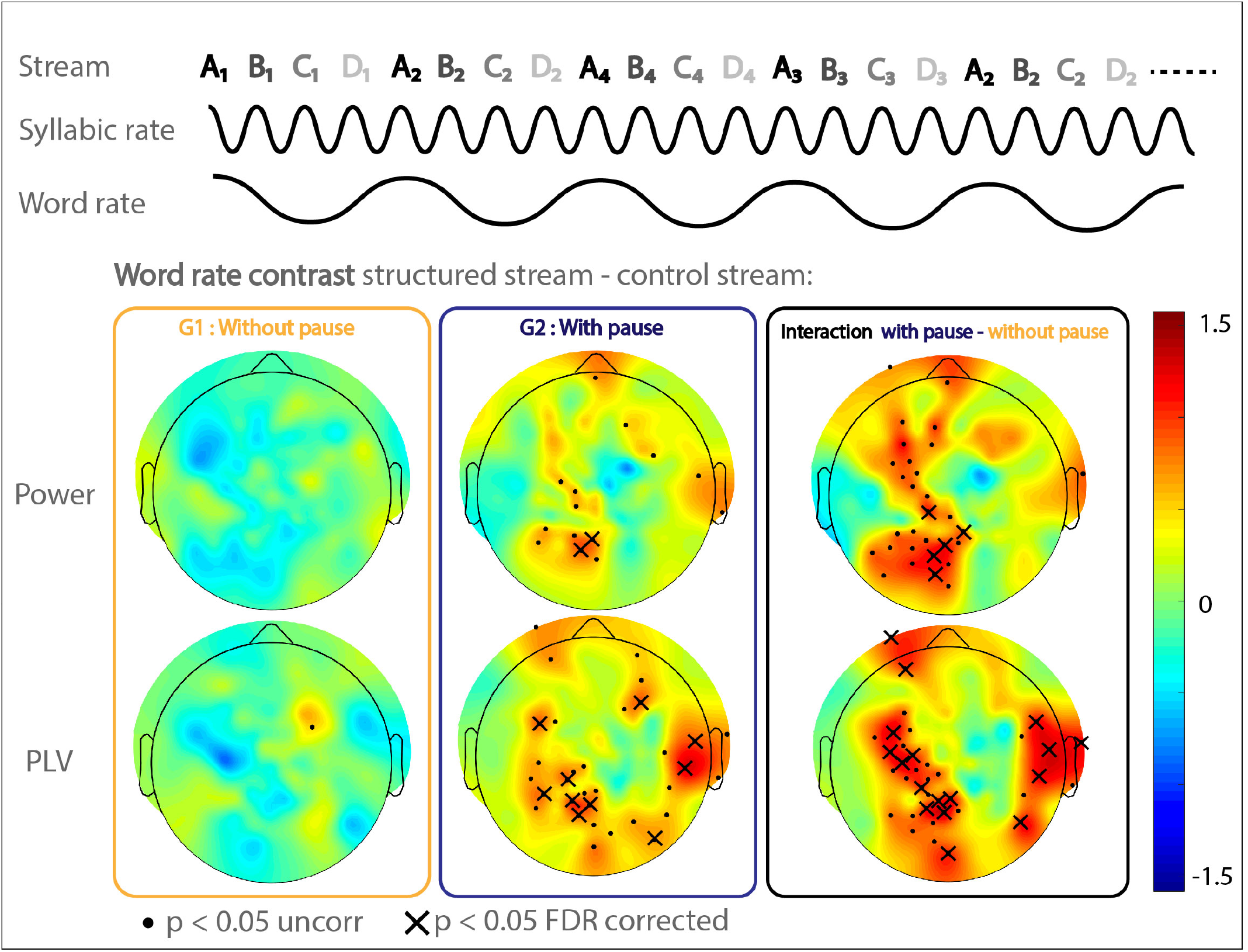
Neural entrainment analyses at the word frequency in the structured stream minus the random stream (Power and phase locking value (PLV)) in the two groups of neonates (G1 and G2). Top rows: the presentation of stimuli with a fixed duration evoked a reproducible time-locked neural response that can be recovered as a neural oscillation at the frequency of stimulation. If infants segment the structure streams based on the quadri-syllabic words, an increase at the word frequency should be observed relative to the random stream. It is what is seen in the second group of neonates (G2) who listened to the streams with sub-liminal pauses. The last column shows the interaction between groups and frequencies. Dots shows the electrodes with a significant result at p<0.05 uncorrected, and x after FDR correction.

It has been described that the power at the syllable rate decreased when adults segment the stream (***Buiatti et al., 2009***). However, we did not find any modulation of the power or PLV at the syllabic frequency in the structured stream compared to the random one.

#### Between-subjects correlation analysis

Because the exact same stream was used in each participant, we were able to analyze whether learning increased neural synchrony between neonates. To do so, we tested whether the correlation between participants increased over time more when they listened to the structured stream. We observed a progressive increase of the mean correlation in neural activity only in G2, exposed to the stream with pauses (Fig 4A Left). Indeed the increase was higher for G2 than G1 (p<0.05, time [564-820]s fig 4 A). During the random streams, the correlations were flat relative to baseline in G1 and G2 (Fig 4B Left). To confirm this effect, we computed a linear regression of the variation of subject correlation with the group with time at the subject level during each stream and compared the slopes in the two groups. Only when neonates listened to the structured stream with pauses (G2), the slope was significantly positive and significantly greater than the same measure in G1 (Fig 4 A Right). No difference was observed during the random streams (Fig 4 B Right).

**Figure 4.**
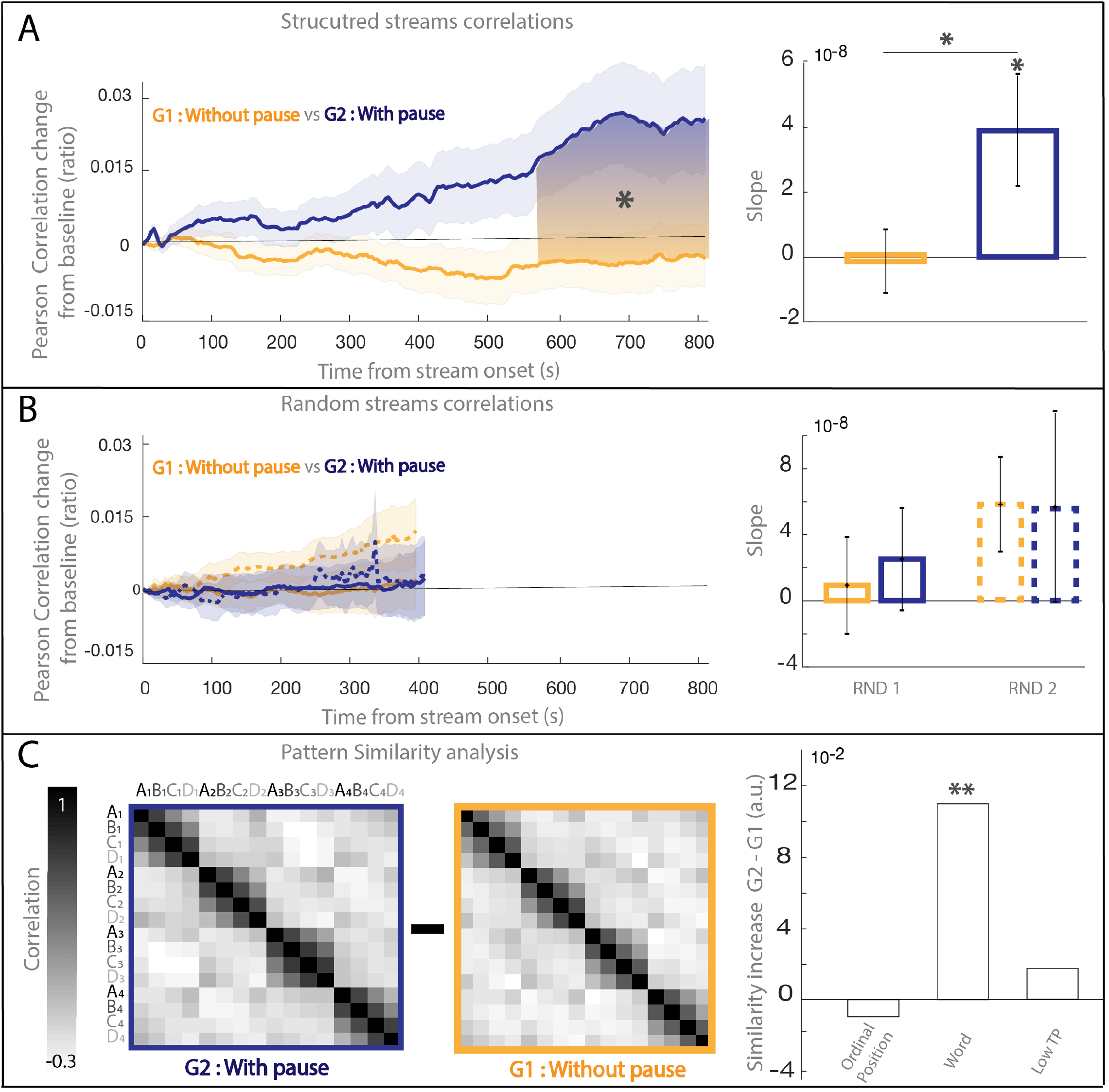
Correlation Analysis A: Comparison of the correlation between neonates in the two groups during the structured stream. Left: Evolution of the correlation across neonates with time. Right: comparison of the slopes of the linear regression with time in each group (yellow G1, blue G2) B: Similar analysis for the first (plain lines) and second random (dotted lines) streams in G1 and G2. C: Pattern Similarity analysis: We computed the increase of pattern similarity between EEG response to each syllables in the two groups during the structured stream. The similarity significantly increased for syllables belonging to the same word (AB, BC, CD, AC, BD, AD)

#### Syllable pattern similarity analysis

In a recent paper, ***Henin et al.*** (***2021***) proposed that pattern similarity between syllables can vary with learning in a similar task. More specifically, using electrocorticography in epileptic adult patients who listened to a structured stream composed of the concatenation of 4 trisyllabic words, they computed different patterns of similarity between the 12 syllables. They observed different clusters of electrodes sensitive either to TP transitions (low vs high TP), the ordinal position (1st vs 2nd vs 3rd syllable), or the word identity (word 1 vs word 2 vs. word 3 vs. word 4). We computed a similar analysis on the responses to the syllables during the structured stream and showed that the similarity pattern for syllables belonging to the same words was significantly increased in G2 (stream with pause) compared to G1 (stream without pause) (p=0.006). However, we failed to find an increase in pattern similarity for low TP (DA’) and for ordinal position (AA, BB, CC and DD) between the two groups. The difference in pattern similarity between the two groups for each condition is reported in fig 4C. To investigate if the significant increase in similarity for syllables belonging to the same word was only due to increase in high TP pairs or in all pairs belonging to the same word, we separated the word condition in two sub-conditions: Consecutive (AB, BC, CD) and non Adjacent (AC, BD & AD). Interestingly, Consecutive and non adjacent pairs showed a significant increase in pattern similarity (both p<0.05) in G2 compared to G1.

#### ERP analysis

Analyses were restricted to regions of interests determined as the two poles of the auditory response to a click preceding the quadri-syllabic element in each group (see method). In G1 (listening to the no-pause stream), there was no significant difference between conditions. Only a trend in the Word vs ShuffleWord comparison was observed in the frontal ROI during two time-windows ([1444-1628]ms, p=0.058 and [1856-1988]ms, p=0.078). Informed by the adult results, we also compared Words and PartWords together against the ShuffleWords - the only condition rejected by the adults. This contrast revealed a significant difference in the frontal ROI (time: [1264-1628]ms, p=0.032)and a trend was observed in the occipital ROI (0.05<p<0.1)(Fig 5 first panel). In G2 (subliminal pauses), ERP to Words differ from those to PartWords and to ShuffleWords both in frontal and occipital ROIs (all p<0.05) (Fig 5 middle panel). (Frontal WvsPW: [1352-1704]ms, Frontal WvsSW: [764-982] & [1080-1304] & [1320-1868]ms, Occipital WvsPW: [1036-1652]ms, Occipital WvsSW [460-1636]ms).

**Figure 5.**
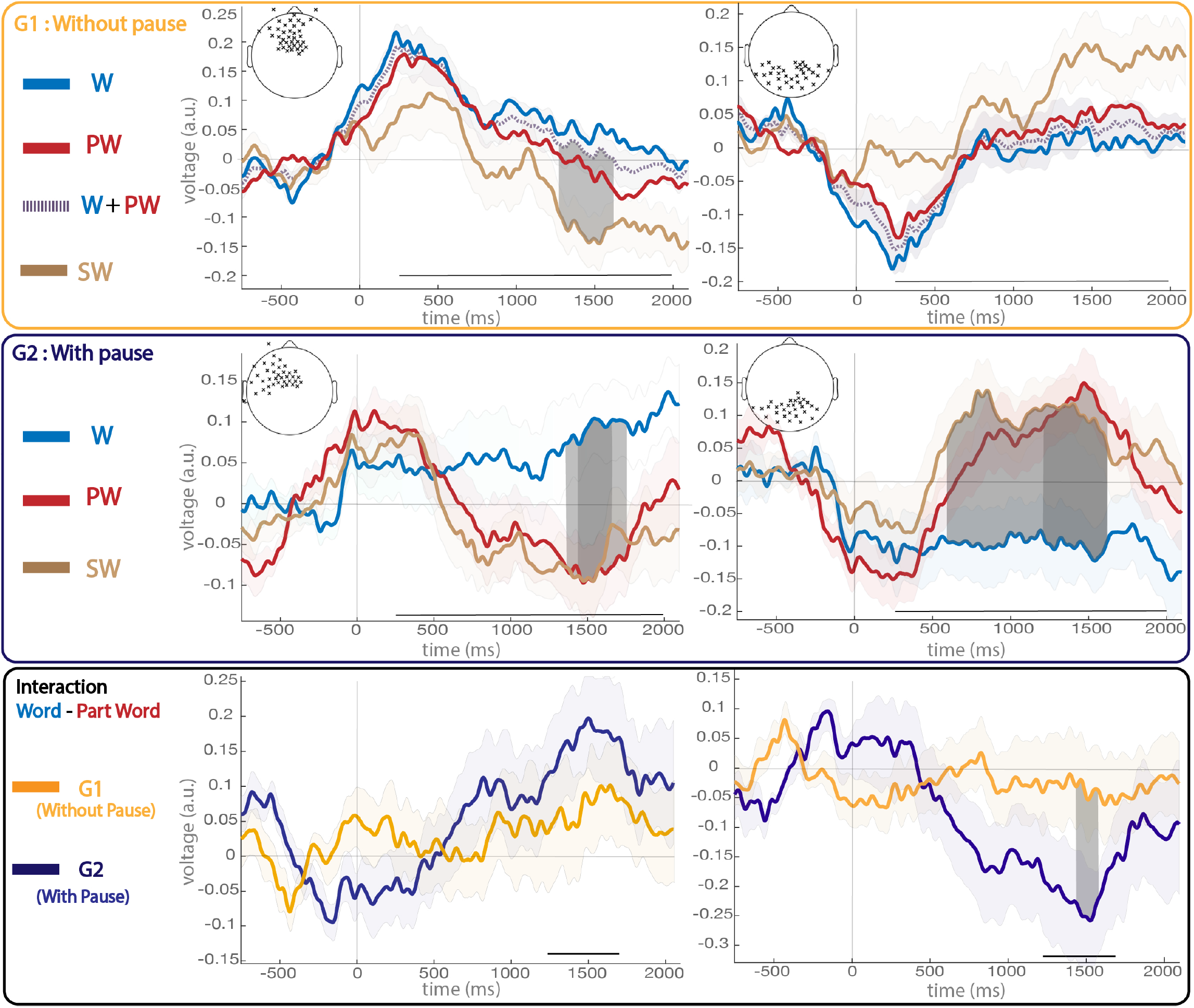
Grand average ERPs to the test-words in G1, G2 and to the word minus part word difference. The ROIs correspond to the two poles of the auditory response to the click preceding the test word in each group. Significant clusters are identified by dark grey areas. Light gray shaded areas surrounding the thick lines represent the standard error across neonates. W=Words (ABCD), PW=Part Words (CDA’B’) and SW=Shuffle Words (ACBD). Gray lines at the bottom of the plots indicate the time windows on which statistics were performed

We also tested the interaction on the main effect Words - Part Words and found a significant interaction (p<0.05) on the occipital ROI (Fig 5 last panel).Using a permutation cluster based method (fieldtrip cluster analysis) between 250 and 2000 ms without previous ROI extractions revealed similar results.

## Discussion

Whereas tri-syllabic words are easily extracted from a flat speech stream using TP between syllables in adults (***Saffran et al., 1996b***), infants (***Saffran et al., 1996a***) and even sleeping neonates (***Fló et al., 2021a***), this single cue was insufficient for quadrisyllabic words even in awake vigilant adults, revealing a clear limitation in sequence processing. Yet, transitional probabilities were computed at both ages: Both adults and neonates rejected ShuffleWords, which contained the exact same syllables as the Words, but in the wrong order. The higher number of syllables to be memorized (16 syllables here vs. 12 for tri-syllabic words) and the larger word size do not by themselves explain this sharp limit since the same material, just with the presence of sub-liminal pauses rescued the word extraction process. When a subliminal pause was present at the end of the Word, we recorded several indicators of learning: First, a significant increase in power and phase locking value was observed at the word frequency in the structured stream relative to the random stream. Second, neural synchrony increased between participants only for the structured stream with pauses, further suggesting that neonates were following a similar learning process in that condition. Finally, ERP to Words and PartWords were significantly different. Similarly, adults ranked Words higher than PartWords. All these markers of successful segmentation were lacking when there was no pause, and the differences between the two stream conditions were significant in infants and in adults. Together, these results suggest, first, thatTP computations are not always sufficient to enable stream segmentation and confirm that prosodic cues are more effective; second, that once the stream is divided into smaller units, memorization is facilitated allowing Words and PartWords to be distinguished. Interestingly, attentive adults were no better than sleeping infants revealing a continuity between ages, and also between vigilance states, thus highlighting a core automatic mechanism at work from birth on. Finally in neonates, the difference between Words and the other conditions was late, i.e. not as early as the first syllable as has been reported for tri-syllabic words (***Fló et al., 2021a***).

### Transition probability tracking does not imply segmenting the speech stream

It was proposed that the computation of the transition probabilities might be used to segment a speech stream, either through boundary markers -a TP drop creating a prediction error and the surprise allowing to memorize the syllable following the drop (i.e. the first syllable of the following word)- or because similarity increases between adjacent events. However, neither the local drop of TP nor the temporal proximity within a chunk were sufficient to structure the stream, even after 13 mins of exposure when the unit size was 4 syllables (i.e. 1s long) whereas in the same circumstances and only after 2 mins of exposure, sleeping neonates perceived a tri-syllabic rhythm and furthermore memorized the set of possible first syllables (***Fló et al., 2021a***). This sharp difference is coherent with the 4±1 limit of the auditory short-term memory (STM) (***Cowan, 2001***), and thus emphasize the role of STM in stream chunking. In the neonates, no rehearsal was possible because they were asleep and, in any case, unable at that age to repeat syllables. Nevertheless, their performance was similar to that of awake competent adults, identifying a structural limitation of the STM. This limit of 4 in STM has been proposed as explaining several linguistic observations, such as the size of phrasal verbs and idioms predominantly used in spoken languages such as English, the mean length of continuous discourse without pauses (***Green, 2017***) and the drop in mutual information scores after four words in many languages (***Pothos and Juola, 2007***). It might thus be possible that the TP drop is only noticeable when the whole word plus the next syllable are present at once in the STM, which is compatible with bi-syllabic and tri-syllabic words but not with quadri-syllabic words. This limit in the size of the chunks explains why TP might not be the most powerful mechanisms to extract words from the speech stream and that other cues should rescue this mechanism.

### Rescuing segmentation with sub-liminal pauses

Adding a subliminal pause at the end of the word radically affects the performances at both ages. Although not consciously perceived, pauses act as other word boundary markers (lengthening of the last syllable, pitch drop, etc..), that neonates can perceive (***Christophe et al., 1994***). Our result demonstrate that such a word boundary marker is not only perceived but is effectively used to segment a stream from birth on, that is before infants have perceived many isolated words. The use of such acoustic cues is then probably not a consequence of language learning but part of the auditory/linguistic perceptive system. We also observed that similarity between syllables within the word was increased relatively to the no-pause stream (Figure 4 C). Not only similarity between adjacent syllables within a word was stronger in the stream with pauses than without pauses but similarity also increased between more distant syllables belonging to the same word. We cannot disentangle whether this increase in similarity between the syllables of a word induced the chunking, or the reverse i.e. because syllables were perceived in the same chunk, their similarity increased. It is also interesting to note that perceiving the stream at a more complex level of representations also increased neural synchrony between the neonates. Whereas the syllabic rate itself, which affects many channels (see Fig2 in appendix), should already create the same entrainment across participants, it is not this low-level cue which is predominant in the neural synchrony between neonates but the perception of a higher level of organization of the stream that allowed more robust responses shared across neonates. Finally, the performances between the test phase during which isolated quadri-syllabic sequences were presented, were also massively affected by the stream condition suggesting that once segmentation was done, memory encoding was improved. Words and PartWords were indeed only discriminated after the stream with pauses. However in a similar experimental paradigm but after a stream of concatenated tri-syllabic words, Words were recognized since the first syllable (***Fló et al., 2021a***) whereas here the difference was developing from around 500 ms to become significant only after the end of the word. The lack of first syllable effect was confirmed by the absence of difference between PartWords and ShuffleWords although the latter were starting with a correct first syllable. It is also consistent with the lack of similarity increase between G1 and G2 within the set of first syllables (Fig4C right), which constrasts with the result reported in adults by Henin et al (2021). Thus, contrary to the tri-syllabic stream, the ordinal position of the syllables was not encoded and the difference between correct and incorrect chunks was longer to develop.

### A sharp distinction between tri and quadri-syllabic words

The differences between the two studies in which the size of the word was changing from 3 to 4 raised interesting questions concerning memory encoding. In ***Fló et al.*** (***2021a***), during the test-phase, infants were no more sensitive to TP (i.e. no distinctive ERP response for triplets containing a TP=0) and were reacting to an incorrect first syllable. Here infants were rejecting ShuffleWords, thus were sensitive to TP, but did not react particularly fast to the incorrect first syllable suggesting a more general response to the global familiarity of the word rather than noticing a particular error. Furthermore, contrary to the increase of similarity between syllables at the same ordinal position with learning reported by Henin et al (2021) in adults, we observed no effect of ordinal position (i.e. here reduced to the first position) in the comparison of the similarity analyses between the two streams. Thus we have no evidence that the first syllable had acquired a particular status contrary to what we observed in the case of tri-syllablic words ***Fló et al.*** (***2021a***). It might be explained by the difference in perception of the drop of TP in a tri-syllabic word stream. The drop which can induce a prediction error for the next syllable and thus a surprise, might favor the encoding of these syllables at a particular position (i.e. the first position of the next word). These results suggest that depending on the chunking cue, different memory processes might be engaged and that TP computation might favor a more precise encoding of the chunking elements, starting with the first syllable. Such an intriguing hypothesis should be further tested in experiments contrasting these two cues and the word-size.

### Similarity between adults and neonate cognitive abilities

In this set of experiments, despite very different measuring methods and attentional state, neonates and adults results were remarkably similar. This is all the more remarkable since, beyond the differences in neural maturation and linguistic expertise, the adults were awake and the neonates asleep. It points to automatic and efficient auditory sequence analyses processes shared across age. Many of the structures that support sequences learning (***Henin et al., 2021***) – hippocampus, dorsal linguistic pathway, the superior temporal region-change rapidly in the first year of life; but the classic assumption that immature means poorly functional is increasingly challenged by brain imaging methods that provide markers of learning in young children. FMRI remains difficult in infants but some results support this hypothesis. In a recent paper (***Ellis et al., 2020***) and colleagues tested 3 to 24 months old babies on a statistical learning task in the visual domain with fMRI and reported activation in the hippocampus associated with chunking. ***Dehaene-Lambertz et al.*** (***2002***) reported activations in temporal and frontal areas in 3-month-olds listening to speech proving activity in the same regions found activated by a similar statistical learning task as here in adults (***Henin et al., 2021***). Note however that sleep in infants is differently organized than later with only two stages, quiet and active sleep. Neonates are awake during short periods, immediately followed by active sleep, which is the equivalent of REM sleep in adults. In adults, learning has been shown to exist during REM (***Andrillon and Kouider, 2016***) and also that a task started during wake can continue during REM (***Andrillon et al., 2016***), opening the possibilities that neonates might process speech, learn and consolidate more efficiently than later on.

### Methodological considerations

Together, our results show that behavioral subjective ranking and EEG analyses provide powerful tools to investigate statistical learning and segmenting tasks. While the behavioral approach seems to give cleaner and statistically more powerful results, there was a great overlap with the neural markers observed in EEG data. Moreover, EEG data enables the investigation of such questions in preverbal and non-verbal subjects with different levels of attention (neonates, sleeping subjects, comatose patients…). Power and PLV during the stream as well as ERP during isolated test words were already proposed as reliable neural markers in this task (***Kabdebon et al., 2015***; ***Fló et al., 2021a***). However, to our knowledge, between subject correlation as a function of time has not been shown to successfully capture some learning behavior in infants yet. Our results confirms that despite noise in infants EEG data, a significant part of the variance cannot be explained by low level bottom up activation to external stimuli but instead by more sustained learning effect. Although this first attempt might have been still noisy, we might hope that this method could more accurately quantify the average amount of learning of a group over time or even characterize learning at the subject level. One drawback of this method is that, to compare across subjects, all subjects have to be exposed to the exact same stimuli, which presents a risk of confound in the experimental design. Here we minimized this risk by taking two precautions. We first carefully designed and balanced the auditory material on acoustic aspects (see Appendix 1). Secondly, we ran two groups with a minimal change (a subliminal pause every four syllables) so that, if any bias persists, it would be the same in both groups and thus cannot explain differences between groups. We also implemented what we believe to be an improvement for ERP analysis. Before the presentation of isolated words, we presented a short audio click. In this way, we were able to extract ROIs for analysis with a data-driven approach. We performed a cluster based-permutation analysis on all data against zero during the click presentation to extract the auditory ERP regions of interest (ROI). Moreover, this cluster was representative of the auditory response in this particular group of subjects taking into account non-relevant variations due to 1) Experimental conditions such as the placement of the net on the infant’s head which is more variable at this age due to birth-related head deformation and can introduce between groups differences 2) eventually long-tail effects of the previous trials on the topography that can affect the baseline.

### Conclusion

Human neonates display sequence learning abilities even during sleep, based on TP computations and segmenting helped by acoustic/prosodic cues. The similarities with adults were remarkable revealing that a speech stream was not a uniform landscape for infants but that different cues might help them to chunk it in smaller units, opening the possibility to discover the linguistic regularities and the productive properties of speech.

## Methods and Materials

### Adult Behavioral Experiment

#### Participants

A total of 43 adults were recruited via social media and mailing (21 males, age distribution = [18-25]: 9, [25-40]:16, [40-60]: 17, 60+: 1]) with no reported auditory issue or language related troubles. They were randomly assigned to one of the streams with the instruction to carefully listen during 3 minutes to a nonsense language composed of nonsense words that they have to learn because they will have to answer questions on the words afterwards. This project was approved by the Ethical research committee of Paris Saclay University under the reference CER-Paris-Saclay-2019-063.

#### Stimuli

All speech stimuli were generated with the MBROLA text-to-speech software (Dutoit, 1997), using French diphones. The duration of all syllables was equalized to 250 ms with flat intonation and no coarticulation between syllables. Each experiment was composed of 3.3 minutes (800 syllables) of an artificial monotonous stream of concatenated syllables, that correspond to the four possible words randomly concatenated with the only restriction that the same word could not be presented twice in a row. The same vocabulary (sixteen syllables) was used in the two streams, with and without pause. In the stream with pause, a 25-ms pause was inserted every 4-syllables. All streams were ramped up and down during the first and last 5 s to avoid the start and end of the streams serving as perceptual anchors. We used the same syllables and words for the infant experiment. To avoid phonological similarity effects that could bias toward one or the other condition, Words and PartWords were reversed for half of the subjects.

In a previous experiment with similar streams with and without 25 ms pauses, ***Peña et al.*** (***2002***) showed that participants were at chance when they had to choose which of the two streams had pauses. To confirm that the pauses were not consciously perceived, 8 adults listened to 20 streams (40 syllables −10 seconds) presented randomly (10 without pauses and 10 with a 25 ms-pause every four syllables) and were unable to indicate which stream had pauses or not (mean = 49% (range [40, 59]%); p = 0.89).

#### Procedure

After listening to the structured stream, participants were asked to rank the familiarity of the individual words (from “Completely unfamiliar” to “Completely familiar”on a 6-step scale). Six conditions (three bi-syllabic and three quadri-syllabic) were used to avoid any bias based on the length of the test words, with 4 trials in each of the 6 conditions. To avoid phonological similarity effects that could bias toward one or the other condition, participants were assigned to one of two groups where conditions were reversed. Four different pairs of structured streams per group were also generated and participants randomly assigned to one pair to avoid any given particularity of a stream driving the results. Three conditions were studied: Words, PartWords and ShuffleWords. Words corresponded to the words that were embedded in the structured streams (ABCD) while PartWords corresponded to the two last syllables of a word and the two first of another word (CDA’B’). Thus, although PartWords were heard during the structured stream, they violated a chunking based on TP. ShuffleWords corresponded to words in which the second and third syllables were inverted, creating a null TP between all syllables (none of the transitions were heard during the structured stream). However, the first and last syllables were correct.

#### Data Processing

Each answer was converted to a numerical value from 1 (completely unfamiliar) to 6 (completely familiar). All data were aggregated together, in each group to compute a linear mixed-effects model on items (y condition + (1|subject)) in order to take the subject effect into account. The p-values were then FDR corrected. To compare subjects’ segmenting performances for both streams, we computed the mean familiarity ranking for each condition in each subject and subtracted the PartWord ranking from the word ranking within each subject. We then performed a one-way unpaired t-test between the two groups.

### Infant EEG Experiment

#### Participants

Two groups of healthy full-term neonates were tested between day 1 and 3. There was no problem during pregnancy and delivery, birth weigth > 2500 g, term > 38 wGA, APGAR ≥ 6 and 8 at 1’ and 5’, normal audition tested with otoacoustic emission. Parents provided informed consent, and the Ethical Committee (CPP Tours Region Centre Ouest 1) approved the study (EudraCT/ID RCB: 2017-A00513-50). In the first group (G1: stream without pause), 34 neonates were tested. Among them 7 were excluded because they did not complete the experimental protocol or because of technical issues leaving 27 infants (14 males). A second group of 34 infants was tested (G2, stream with pauses); 9 were excluded because they did not complete the experimental protocol or because of technical issues, leaving 25 infants (13 males).

#### Stimuli

We used the same 16 isolated syllables generated with MBROLA as in the adult experiment to construct 4 different streams (structured and random, with and without pause). The random stream was around 6.5 mn-long and consisted of 1600 pseudo-randomly concatenated syllables. Each syllable could be followed by three others from the pool leading to a flat TP during the stream. This pseudo-random stream offers a more controlled stimulus than the random streams used previously because the TP were fixed to (instead of 1/15), that is a similar value than the TP between words in the structured stream. The structured stream was about 13 mn-long and comprised 3200 syllables. All streams were ramped up and down during the first and last 5 s to prevent the beginning and the end of the streams from being used as anchors. We created only one syllabic order for each stream with the goal to obtain learning markers comparable between infants. For G2, a pause was added every 4 syllables in both the structured and the random streams. Thus the sequences were identical for all infants in the two groups except for the 25-ms subliminal pauses every 4 syllables in the second group. Because all subjects had the same auditory materials, we carefully controlled for low level acoustic-phonetic properties. We equilibrated the characteristics of consonants and vowels in the different words and at the different syllabic positions within words to avoid that learning be based on low-level acoustic cues (See fig S1 for more details). As in adults, three types of test words were created: Word (ABCD), PartWord (CBA’B’) and ShuffleWords (ACBD). (Table 1 and Fig S1).

**Table 1.** List of words used for the infant experiment. During each small test block, 12 test words were presented in isolation, 4 Words, 4 ShuffleWords and 4 PartWords out of the 8 possible.

| Words (W) | Part Words (PW) | Shuffle Words (SW) |
| --- | --- | --- |
| RaFiBouNeu | BouNeuNonLo / BouNeuVouDon | RaBouFiNeu |
| GuMaReuZo | ReuZoVouDon / ReZoNonLo | GuReuMaZo |
| NonLoSanBi | SanBiGuMa / SanBiRaFi | NonSanLoBi |
| VouDonMuLan | MuLanRaFi / MuLanGuMa | VouMuDonLan |

#### Procedure

EEG was recorded with 128 electrodes (EGI geodesic sensor net), carefully placed on the neonates’ head by trained researchers to increase the consistency of the net placement. 3 nets with different radii were used to fit infants’ head. For G1, infants were tested while asleep in the experimenter or parent’s arms. Due to COVID restrictions, the second group of babies was tested asleep in the crib. Both groups followed the same procedure (Fig 1). A first control stream of pseudorandom concatenation of 1600 syllables was followed by a structured stream composed of 3200 syllables grouped in words of 4 syllables. Infants then heard eight repetitions of short structured streams (160 syllables) followed by 12 test words presented in isolation (4 in each condition: Word, PartWords and ShuffleWords, ISI 2-2.5s) for a total of 96 test-words (32 in each condition). The short streams were added to maintain learning because 2/3 of the test-words were violating the learned structure. Each test word was preceded by a short click 200 ms before its onset. This was added to reset the baseline with a neutral event to avoid long-range effects following the words and thus affecting the following stimulus and also to extract auditory ROI from task unrelated auditory stimuli. Finally, a second control stream was presented. Thus the two random-streams were sandwiching the structured stream in order to control for habituation to the auditory stimulation, change in sleep stage and any confounding time effect.

#### Data Processing

EEG recordings were band-pass filtered between 0.2 and 15Hz for all analyses. Artifact rejection was performed on the non-epoched recording session using APICE pipeline (***Fló et al., 2021b***) based on the EEGLAB toolbox (***Delorme and Makeig, 2004***). Artifacts were identified on continuous data, based on voltage amplitude, variance, first derivative, and running average. The variance algorithm was applied in sliding time windows of 500 ms with 100 ms steps. Adaptive thresholds were established for each subject and electrode as two interquartile ranges away from the 3rd quartile. This gave as result a logical matrix ofthe size ofthe data indicating bad data. Electrodes were definitely rejected if they were marked as bad more than 50% of the recording time, and time-samples were marked as bad if more than 35% of the electrodes were marked bad at this time-sample. For the ERP analysis, we then performed spatial interpolation of missing channels and the data were mathematically referenced to the average of the 128 channels. Due to COVID protocols, experimental setups were slightly different for the groups. Human arms are more adapted to support neonates than cribs, thus there were slightly more artifacts in the second group. To avoid too much rejection for the second group, we slightly decreased the stringency of the artifact rejection algorithms in the second group (mean rejection rate for G1: 15.7% (std 6.3%) vs. 25.9% for G2 (std 13.4%).

#### Neural entrainment

The recordings from the structured stream and the random streams were segmented into consecutive non-overlapping epochs of 15 seconds (corresponding to 15 quadrisyllabic words in the first experiment and 14.6 due to the pauses in the second). All subjects having more than 6 good epochs in each condition were included in this analysis (27 neonates in G1, 25 in G2). For each neonate and electrode, we averaged the activity over artifact-free epochs and computed the Fourier Transform using the fast Fourier transform algorithm (FFT) as implemented in MATLAB. We then computed the power of the FFT. The Phase Locking value (PLV) between trials was computed on the FFT of single trials. The frequencies of interest were selected as the inverse of the duration of a word (f = 1Hz for the first group f=0.98 for the second) and one-quarter of a word (i.e. roughly a syllabic rate, f = 4 Hz for the first group, f = 3.9 Hz for the second). To assess significance of the power/PLV at the two frequencies of interest, we computed a contrast between the power/PLV during the structured stream compared to the random streams for each electrode. As we expect learning during the structured stream to elicit a word rate oscillation, we computed a one-way (structured>random) paired t-test on each electrode. The p-values were then FDR corrected across electrodes. To look for a potential difference between groups, we then computed an interaction between the previously described contrasts of both groups and ran a one-way unpaired t-test on each electrode with a FDR correction for multiple comparison across electrodes.

#### Correlation Analysis

In both experiments, all subjects heard the exact same auditory material avoiding differences in stimulation between participants. We could thus compute instantaneous correlation between each participant and all the others. For each subject at each time during the streams, we computed the correlation at the topographical level between the topography of subject i at time t and the topography of the grand average excluding subject i at time t. It gave for each subject a vector of correlation between its own topography and the mean topography of all other subjects throughout time. Bad data were replaced by zeros and not taken into account for the average topographies across subjects. Time points with only bad data gave NaN correlation results. Our hypothesis was that learning should lead to an increase with time in the correlation between neonates as they learn the same material. To test it, we used two different methods. In the first one, we smoothed the correlation signal using a 400s-sliding-average-window in each neonate and stream, then computed a cluster-based analysis to reveal a significant cluster of time during which one stream showed a greater correlation than the other one. In the second one, we computed the slope of the linear regression with time in each subject, and then considered the slope as variable for the structured and random conditions in t-test comparing both groups.

#### Pattern Similarity Analysis

To compute pattern similarity between syllables, we epoched each syllable from the structured stream from −100 ms to 350 ms. We removed the 100 first syllables to give enough time for participants to learn the task. The remaining epochs were averaged by syllables for each subject and a correlation matrix between each pair of syllables was computed with all the electrodes between 0 and 350 ms. We then separated the pairs of syllables in 5 conditions: First syllable (AA), Ordinal position (BB or CC or DD), Word and TP (AB or BC or CD), Word only (AC or AD or BD) and Low TP (DA’). We then averaged the similarity per condition and subtracted the correlation between all the other pairs. We could then compare between the two groups if pattern similarity between groups of syllables was increased by learning how to segment the stream (One-way t-test G2>G1).

#### ERP Analysis

Data were segmented in 2850 ms long epochs ([−750 +2100]ms relative to word onset), averaged in the three conditions (Words, PartWords and ShuffleWords) and baseline-corrected with the mean voltage value in the interval [−750 - 0]. Neonates with less than 5 remaining trials in one of the three conditions were excluded from analysis (1 neonate in G1 and 1 in G2) To identify regions of interest adapted to each group, we took advantage of the click presentation to extract the auditory event-related potential associated with auditory response by running a cluster-based analysis (5000 randomizations, two tailed t-test, alpha < 0.01, cluster-alpha < 0.01) between −200 and 0 ms. This procedure identifies in each group a positive frontal and a negative occipital clusters, on which we restricted the ERP analyses. In each neonate and condition, the voltage was averaged across electrodes in each cluster. A cluster-based analysis was performed on the time-series (5000 randomizations two tailed t-test alpha < 0.05, cluster alpha < 0.05) between 250ms (end of the first syllable) and 2000 ms to compare all pairs of conditions. Because of the adults’ behavioral results, we added the contrast ‘heard’ (average of Word and PartWord) vs ‘non heard’ (ShuffleWord) in G1. Finally, we computed the interaction between groups and conditions (Word-PartWord) during the time-window in which the previous analysis revealed a significant effect.

## Acknowledgments

We thank Simon Henin for help and comments about pattern similarity analysis. This research has received funding from the European Research Council (ERC) under the European Union’s Horizon 2020 research and innovation program (grant agreement No. 695710).

## Appendix 1

### Word Design Constraints

In order to perform between subject correlation as a measure of learning during the stream, we exposed all infant participants to the exact same auditory material in the same order. This choice is at risk of bias if by any lack of chance, there is some very particular structure in the streams or any acoustical bias in the words we used. To minimize the risk, we used several methods. First, both groups (with and without pauses) had the same auditory material in the same order. The only difference was the added pauses for the second group. Thus any bias should have similarly affected the two groups. Secondly, we carefully designed the auditory material that was presented to the participants to avoid any particular low-level acoustical feature to drive the effect. Finally, we ran the same experiment with adults with a more randomized approach between participants and replicated the results. Altogether, this great care in the stimuli design allowed us to perform a new between-subjects analysis and find a new neural correlate of learning in infant EEG data.

The balance of the low level acoustical feature is presented in fig S1: we applied many rules so that consonants and vowels with different acoustical properties were balanced between words.

**Appendix 1 Figure 1.**
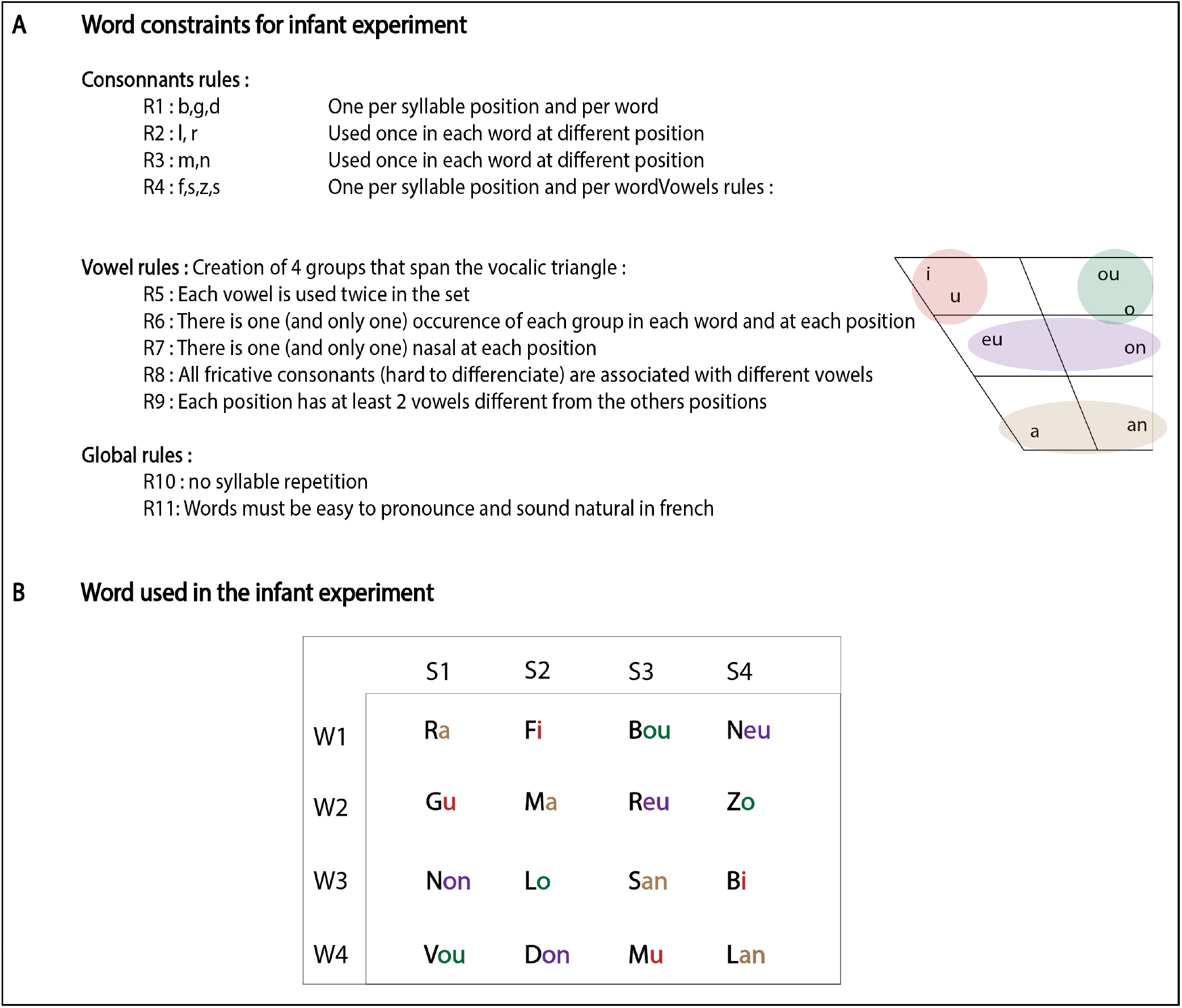
A: List of constraints applied to create an unbiased set of word for the neonate experiment; B: Words used for the neonate experiment

### Neural entrainment power

As described in the main text, we computed the power of the neural entrainment at the syllabic word frequencies for both structured and random streams. In the main figure (fig 3) we only presented the contrast at the word rate between the structured and random streams. We showed a significant increase of the word rate power during the structured stream only G2. For the sake of completeness, we present here the results of the power at word and syllabic rates, for structured and random streams for both groups. For each frequency of interest, we estimated the significance of the power for each electrode by comparing to the power of neighbour frequencies.

**Appendix 1 Figure 2.**
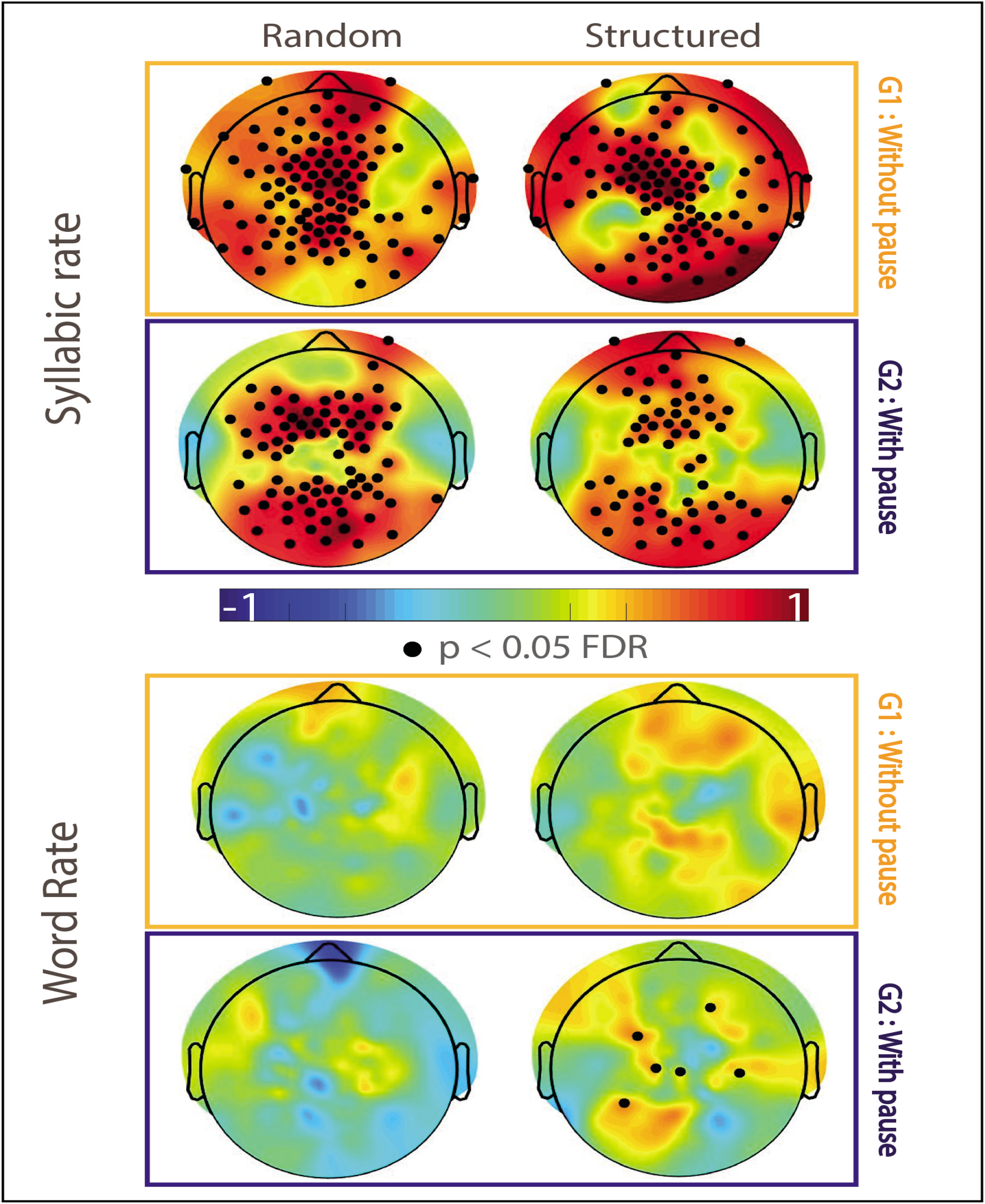
Power ofthe neural entrainment at syllabic (top) and word (botton) rates for random (left) and structured (right) stream in G1 (orange) and G2 (blue)

## References

Andrillon T, Kouider S. Implicit memory for words heard during sleep. Neuroscience of Consciousness. 2016; 2016(1):niw014. doi: 10.1093/nc/niw014.

Andrillon T, Poulsen AT, Hansen LK, LÉ Ger D, Kouider S. Neural markers of responsiveness to the environment in human sleep. Journal of Neuroscience. 2016; 36(24):6583–6596. doi: 10.1523/JNEUROSCI.0902-16.2016.

Bagou O, Frauenfelder UH. Lexical Segmentation in Artificial Word Learning: The Effects of Converging Sublexical Cues. Language and Speech. 2018; 61(1):3–30. doi: 10.1177/0023830917694664.

Batterink LJ, Choi D. Optimizing steady-state responses to index statistical learning: Response to Benjamin and colleagues. Cortex. 2021; https://doi.org/10.1016/j.cortex.2021.06.008., doi: 10.1016/j.cortex.2021.06.008.

Benjamin L, Dehaene-Lambertz G, Fló A. Remarks on the analysis of steady-state responses: spurious artifacts introduced by overlapping epochs. Cortex. 2021;https://doi.org/10.1016/j.cortex.2021.05.023, doi: 10.1016/j.cortex.2021.05.023.

Black A, Bergmann C. Quantifying infants’ statistical word segmentation: A meta-analysis. Proceedings of the 39th Annual Conference of the Cognitive Science Society. 2017; (3):124–129. https://pdfs.semanticscholar.org/0807/41051b6e2b74d2a1fc2e568c3dd11224984b.pdf.

Buiatti M, Peña M, Dehaene-Lambertz G. Investigating the neural correlates of continuous speech computation with frequency-tagged neuroelectric responses. NeuroImage. 2009; 44(2):509–519. http://dx.doi.org/10.1016/j.neuroimage.2008.09.015, doi: 10.1016/j.neuroimage.2008.09.015.

Christophe A, Dupoux E, Bertoncini J, Mehler J. Do infants perceive word boundaries? An empirical study ofthe bootstrapping of lexical acquisition. Journal of the Acoustical Society of America. 1994; 95(3):1570–1580. doi: 10.1121/1.408544.

Cowan N. The magical number 4 in short-term memory: A reconsideration of mental storage capacity. Behavioral and Brain Sciences. 2001; 24(1):87–114. doi: 10.1017/S0140525X01003922.

Dehaene-Lambertz G, Dehaene S, Hertz-Pannier L. Functional neuroimaging of speech perception in infants. Science. 2002; 298(5600):2013–2015. doi: 10.1126/science.1077066.

Delorme A, Makeig S. EEGLAB: An open source toolbox for analysis of single-trial EEG dynamics including independent component analysis. Journal of Neuroscience Methods. 2004; 134(1):9–21. doi: 10.1016/j.jneumeth.2003.10.009.

Ellis CT, Skalaban LJ, Yates TS, Bejjanki VR, Córdova NI, Turk-Browne NB. Evidence of hippocampal learning in human infants. bioRxiv. 2020; doi: 10.1101/2020.10.07.329862.

Ferry AL, Fló A, Brusini P, Cattarossi L, Macagno F, Nespor M, Mehler J. On the edge of language acquisition: Inherent constraints on encoding multisyllabic sequences in the neonate brain. Developmental Science. 2016; 19(3):488–503. doi: 10.1111/desc.12323.

Fiser J, Aslin RN. Statistical learning of new visual feature combinations by infants. Proceedings of the National Academy of Sciences of the United States of America. 2002; 99(24):15822–15826. doi: 10.1073/pnas.232472899.

Fló A, Benjamin L, Palu M, Dehane-lambertz G. From computing transition probabilities to word recognition in sleeping neonates, a two-step neural tale. bioRxiv. 2021; p. 1–26.

Fló A, Brusini P, Macagno F, Nespor M, Mehler J, Ferry AL. Newborns are sensitive to multiple cues for word segmentation in continuous speech. Developmental Science. 2019; 22(4). doi: 10.1111/desc.12802.

Fló A, Gennari G, Benjamin L, Dehaene-lambertz G. Automated Pipeline for Infants Continuous EEG (APICE): a flexible pipeline for developmental studies. bioRxiv. 2021;.

Green C. Usage-based linguistics and the magic number four. Cognitive Linguistics. 2017; 28(2):209–237. doi: 10.1515/cog-2015-0112.

Hauser MD, Newport EL, Aslin RN. Segmentation of the speech stream in a non-human primate: Statistical learning in cotton-top tamarins. Cognition. 2001; 78(3):53–64. doi: 10.1016/S0010-0277(00)00132-3.

Henin S, Turk-Browne NB, Friedman D, Liu A, Dugan P, Flinker A, Doyle W, Devinsky O, Melloni L. Learning hierarchical sequence representations across human cortex and hippocampus. Science Advances. 2021; 7(8):1–13. doi: 10.1126/sciadv.abc4530.

Hochmann JR, Endress AD, Mehler J. Word frequency as a cue for identifying function words in infancy. Cognition. 2010; 115(3):444–457. http://dx.doi.org/10.1016/j.cognition.2010.03.006, doi: 10.1016/j.cognition.2010.03.006.

James LS, Sun H, Wada K, Sakata JT. Statistical learning for vocal sequence acquisition in a songbird. Scientific Reports. 2020; 10(1):1–18. doi: 10.1038/s41598-020-58983-8.

Johnson EK, Tyler MD. Testing the limits of statistical learning for word segmentation. Developmental Science. 2010; 13(2):339–345. doi: 10.1111/j.1467-7687.2009.00886.x.

Kabdebon C, Pena M, Buiatti M, Dehaene-Lambertz G. Electrophysiological evidence of statistical learning of long-distance dependencies in 8-month-old preterm and full-term infants. Brain and Language. 2015; 148:25–36.http://dx.doi.org/10.1016/j.bandl.2015.03.005, doi: 10.1016/j.bandl.2015.03.005.

Mandel DR, Jusczyk PW, Kemler Nelson DG. Does sentential prosody help infants organize and remember speech information? Cognition. 1994; 53(2):155–180. doi: 10.1016/0010-0277(94)90069-8.

Nespor M, Vogel I, Prosodic phonology; 2006. doi: 10.4324/9780203993880-15.

Ordin M, Polyanskaya L, Laka I, Nespor M. Cross-linguistic differences in the use of durational cues for the segmentation of a novel language. Memory and Cognition. 2017; 45(5):863–876. doi: 10.3758/s13421-017-0700-9.

Peña M, Bonatti LL, Nespor M, Mehler J. Signal-driven computations in speech processing. Science. 2002; 298(5593):604–607. doi: 10.1126/sci-ence.1072901.

Pothos EM, Juola P. Characterizing linguistic structure with mutual information. British Journal of Psychology. 2007; 98(2):291–304. doi: 10.1348/000712606X122760.

Saffran JR, Aslin RN, Newport EL. Statistical Learning by 8-Month-Old Infants. Science. 1996; 274(5294):1926–1928.http://www.ncbi.nlm.nih.gov/pubmed/8943209.

Saffran JR, Johnson EK, Aslin RN, Newport EL. Statistical learning of tone sequences by human infants and adults. Cognition. 1999; 70(1):27–52. doi: 10.1016/S0010-0277(98)00075-4.

Saffran JR, Newport EL, Aslin RN. Word segmentation: The role of distributional cues. Journal of Memory and Language. 1996; 35(4):606–621. doi: 10.1006/jmla.1996.0032.

Schön D, Boyer M, Moreno S, Besson M, Peretz I, Kolinsky R. Songs as an aid for language acquisition. Cognition. 2008; 106(2):975–983. doi: 10.1016/j.cognition.2007.03.005.

Shukla M, Nespor M, Mehler J. An interaction between prosody and statistics in the segmentation of fluent speech. Cognitive Psychology. 2007; 54(1):1–32. doi: 10.1016/j.cogpsych.2006.04.002.

Shukla M, White KS, Aslin RN. Prosody guides the rapid mapping of auditory word forms onto visual objects in 6-mo-old infants. Proceedings of the National Academy of Sciences of the United States of America. 2011; 108(15):6038–6043. doi: 10.1073/pnas.1017617108.

Teinonen T, Fellman V, Näätänen R, Alku P, Huotilainen M. Statistical language learning in neonates revealed by event-related brain potentials. BMC Neuroscience. 2009; 10. doi: 10.1186/1471-2202-10-21.

Toro JM, Sinnett S, Soto-Faraco S. Speech segmentation by statistical learning depends on attention. Cognition. 2005; 97(2):25–34. doi: 10.1016/j.cognition.2005.01.006.

Toro JM, Trobalón JB. Statistical computations over a speech stream in a rodent. Perception and Psychophysics. 2005; 67(5):867–875. doi: 10.3758/BF03193539.

Tyler MD, Cutler A. Cross-language differences in cue use for speech segmentation. The Journal of the Acoustical Society of America. 2009; 126(1):367–376. doi: 10.1121/1.3129127.

Wakefield JA, Doughtie EB, Lee Yom BH. The identification of structural components of an unknown language. Journal of Psycholinguistic Research. 1974; 3(3):261–269. doi: 10.1007/BF01069242.

